# How structural elements added by evolution from bacterial transporters to human SLC6 homologs have enabled new functional properties

**DOI:** 10.1101/204164

**Authors:** Asghar M. Razavi, George Khelashvili, Harel Weinstein

## Abstract

Much of the structure-based mechanistic understandings of the function of SLC6A neurotransmitter transporters emerged from the study of their bacterial LeuT-fold homologs. It has become evident, however, that structural differences such as the long N- and C-termini of the eukaryotic neurotransmitter transporters impart an expanded set of functional properties to the eukaryotic transporters, which are not shared by the bacterial homologs that lack the structural elements that appeared later in evolution. However, mechanistic insights into some of the measured functional properties of the eukaryotic transporters, that have been suggested to involve these structural elements, are sparse. To learn how the structural elements added in evolution enable mechanisms of the eukaryotic transporters in ways not shared with their bacterial LeuT-like homologs, we focused on the human dopamine transporter (hDAT) as a prototype. We present the results of a study employing large-scale molecular dynamics simulations and comparative Markov State Model analysis of experimentally determined properties of the wild type and mutant hDAT constructs, which reveal a rich spectrum of interactions of the hDAT N-terminus and the mechanisms by which these contribute to regulation (e.g., by phosphorylation), or to entirely new phenotypes (e.g., reverse uptake – efflux) added in evolution. We reveal separate roles for the distal and proximal segments of the much larger N-terminus shared by the eukaryotic transporters compared to the bacterial ones, consistent with the proposal that the size of this region increased during evolution to enable more, and different, modes of regulation that are not shared with the bacterial homologs.

## Background

The dopamine transporter (DAT) is a member of the neurotransmitter:sodium symporter (NSS) family of proteins belonging to the solute carrier 6 (SLC6) family that performs the reuptake of dopamine from the synaptic cleft into the presynaptic nerve required for neuronal signaling (1). The essential role of DAT, and of its closely related homologs – the serotonin and norepinephrine transporters (SERT and NET) – in signal termination at the synapse makes them important targets for psychostimulants such as cocaine and amphetamines, as well as for pharmaceutical treatment of a variety of disorders of the nervous system (2). Moreover, genetic modifications of the functions of these transporters (3,4) have been implicated in diseases including Schizophrenia, Parkinson’s, and Attention-deficit/hyperactivity disorder (ADHD). Reverse transport of the neurotransmitters (efflux) mediated by DAT and SERT, which has been shown to be affected by such disease-related mutations, is currently a very active topic of research on mechanisms of these membrane proteins (4-15).

Much has been learned about these mammalian neurotransmitter transporters from the investigation of structure and function of their bacterial homologs with which they share many structural and mechanistic properties (16-19). It has become clear, however, that important structural differences exist between the eukaryotic and bacterial proteins, the largest being the much longer N- and C-termini that have been proposed to be partially structured (20,21). Notably, experimental data point to an involvement of these regions of structural difference in measured functional properties of the mammalian NSS (22-24). For example, the phosphorylation of the N-terminus has been implicated the efflux functions of the human DAT (hDAT) (22,25), and our work has shown that the amphetamine (AMPH)-induced reverse transport (efflux) exhibited by DAT and SERT, but not by the bacterial analogs, is dependent on electrostatic interactions between the hDAT N-terminus and negatively charged phosphatidylinositol 4,5-biphosphate (PIP_2_) lipids in the membrane (29,36). Notably, the measured substrate uptake by DAT, a function it shares with the bacterial transporters, is not affected by this N-terminus interaction (35).

Structure-based mechanistic details of the mode in which eukaryotic NSS function is modulated by the involvement of the N-terminus are still sparse, but mechanistic insight from computational studies of hDAT has shown (26) that the N-terminus of DAT engages the PIP_2_ lipid component of membranes to achieve conformational changes related to function (CCRF). These include (i)-the transition of the DAT from outward-facing to inward-facing configurations (27), and (ii)-the release of the sodium ion from the Na2 binding site that is known to precede substrate transport (28,29) and can serves as a monitor of the initial stages of the functional mechanism (30). The complexity of the different CCRF suggested that the N-terminus/PIP_2_ interactions supporting them may (1)-involve different modes of association with the membrane under various conditions (such as in the presence/absence of PIP_2_), and (2)-would be different for different hDAT constructs (e.g., when the N-terminus is phosphorylated or mutated). This reasoning provided a specific testable hypothesis about the mechanism by which new properties of the eukaryotic transporters, those that are not shared with the bacterial homologs, are enabled by the addition of the long N- and C-terminals.

To probe this hypothesis, and verify the relation between the experimental measurements and the specific modes in which the N-terminus participates, we undertook the present computational study of the modes of interaction of the N-terminus in hDAT with the rest of the structure, including the C-terminus, under various conditions and with modifications (phosphorylation, mutations) that have known functional consequences. Here we describe the results from extensive ensemble-level all-atom molecular dynamics simulations we used in this study that also included Markov State Model analysis of hDAT dynamics modulated by mutated and/or modified N-terminus constructs and conditions. The results reveal preferred modes of interaction of the N-terminus with the intracellular domains of hDAT, which can be directly associated with experimentally measured functional phenotypes of the transporter. We show how these interaction patterns change under conditions that have been demonstrated to selectively affect efflux but not regular transport – e.g., PIP_2_ depletion, mutations such as R51W, the K3A/K5A double mutation, or phosphomimic substitution, S/D, of Serine residues at positions 2, 4, 7, 12, and 13 to Aspartate (22,31,32). Moreover, we verify, for the first time, the consistency of observations relating quantitative measures of the specific modes of interaction of the N-terminus with the measured functional properties attributed to them.

The coherent and direct relation between experimentally determined effects of the mutations and conditions, and the interaction modes identified from the simulations, validates the computational results and mechanistic conclusions. Moreover, because the mechanistic inferences are described in atomistic detail, they offer specific experimentally testable predictions for further studies of SLC6 transporter function and of the structure-based relation between the function of bacterial and eukaryotic members of this family. In particular, the details of the rich spectrum of modes of interaction of the long N-terminus of hDAT that emerges from these studies reveal the different roles of the *distal* and *proximal* segments of the N-terminus in modulating specific functions of hDAT. As these are segments of the much larger N-terminus shared by the eukaryotic transporters compared to the bacterial ones, the findings bring mechanistic support for our proposal that the size of this region increased during evolution so as to enable more, and different, modes of regulation that are not shared with the bacterial analogs. An example discussed in detail is the mechanistic explanation for experimentally determined differences in the effects on uptake versus efflux resulting from manipulation of the N-terminus by partial truncation, mutations, and/or elimination of PIP_2_ interactions. This example further underscores the central role of this structural addition in the evolution from the bacterial LeuT-like members of this family.

## Results

The full complement of MD simulation trajectories carried out specifically for this study as described in Methods, includes 50 statistically independent ~1 μs-long trajectories for each of the four different conditions and constructs for which quantitative measurements of activity are available. As discussed before (30) the ensemble exploration of the configurational space of each construct/condition bolsters the statistical validity of the inferences and predictions from the simulation. The specific molecular systems investigated in this manner include: (i)-wild type hDAT in PIP_2_-depleted membranes (hereafter referred to as “no-PIP_2_ system”), (ii)-hDAT with the R51W mutation in the N-terminal domain, with the double K3A+K5A mutation in the N-terminus (termed “K3/5A system”), and with the first five N-terminal Ser residues substituted by Asp as a phosphomimic (termed “S/D construct”); these 3 mutant constructs were immersed in PIP_2_-containing bilayers, consistent with the experimental conditions under which their functional properties were assayed. The data from these computations are compared to results for the wild type hDAT simulated in PIP_2_-containing membranes we reported recently (30), and are analyzed utilizing the same protocols as described therein and detailed here in Methods.

### Different modes of interaction of the N-terminus with the rest of the hDAT protein correspond to differences in experimentally measured functional properties

From the simulation trajectories we identified the regions of hDAT structure that interact with the N-terminus, and generated the per-residue contact map shown in Figure 1. This map shows that with PIP_2_ present in the membrane, the N-terminus, as a whole, interacts with all intracellular loop (IL) regions of hDAT (IL1, IL2, IL3, IL4, and IL5), and with the C-terminus.

**Figure 1.**
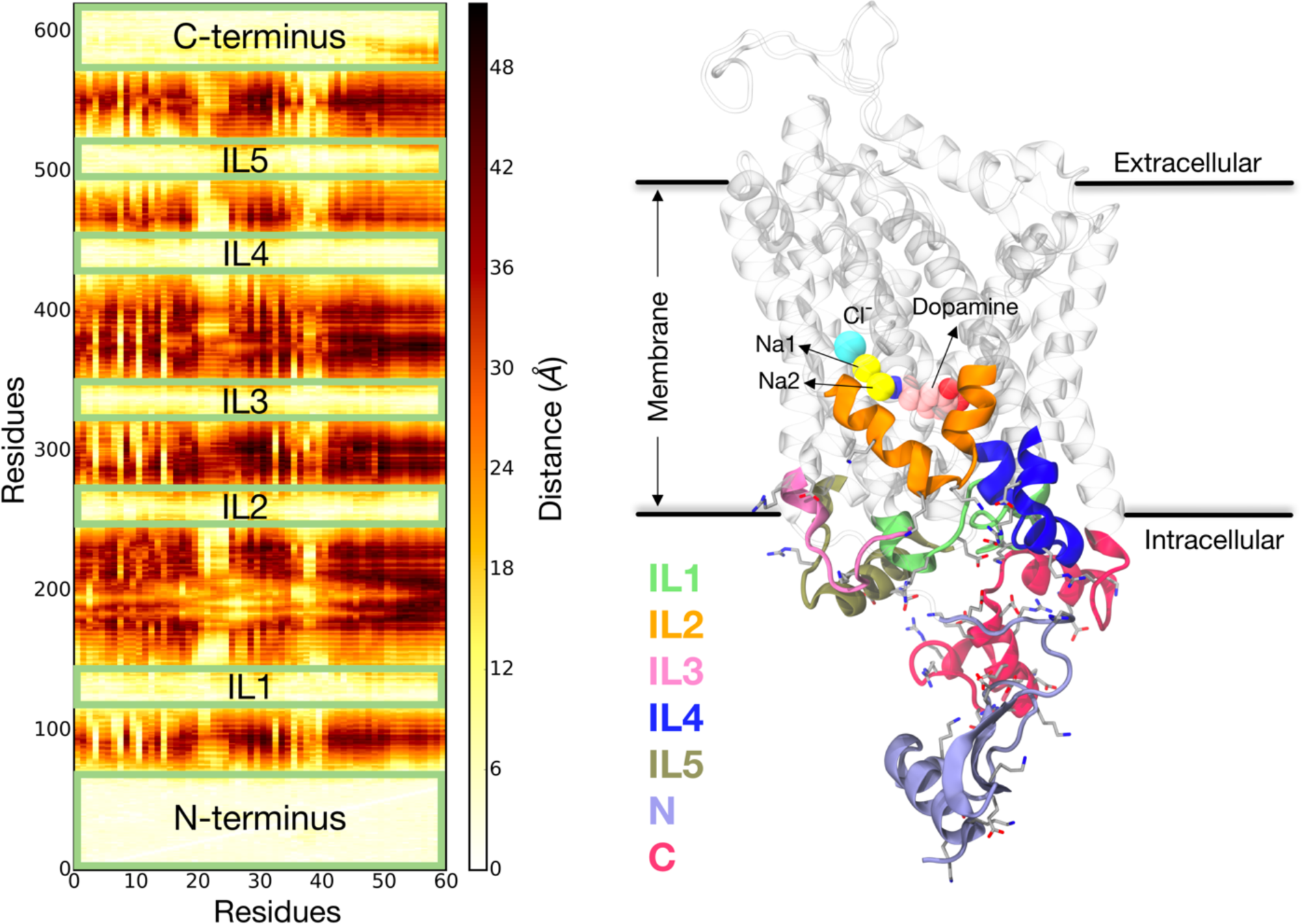
hDAT structure and N-terminus interactions. (**Left**) Contact map for interaction of N-terminus residues (x-axis) with all hDAT residues (y-axis). Distance calculations were done with the *closest-heavy algorithm* implemented in MDTraj software (33). The color bar shows the minimum distance of each residue in the N-terminus to other residues (the lightest colored regions indicate the strongest interactions, e.g., in the various loop segments) as identified in the ensemble of 50 μs trajectories for the wild-type hDAT embedded in the PIP_2_-containing lipid bilayer (see Methods). (**Right**) Snapshot of hDAT structure highlighting the intracellular segments interacting with the N-terminus in the ensemble of 50 μs trajectories. Charged residues are shown in licorice.

A detailed comparative analysis of the interactions between the N-terminus with the intracellular regions of wild type hDAT in PIP_2_-containing membranes (obtained from equivalent trajectories described recently (30)) and the constructs studied here (including PIP_2_-depleted membrane conditions) reveals a specific pattern (modes) of interaction of the different parts of the N-terminus with intracellular regions of the transporter. These patterns are presented in Figure 2.

**Figure 2.**
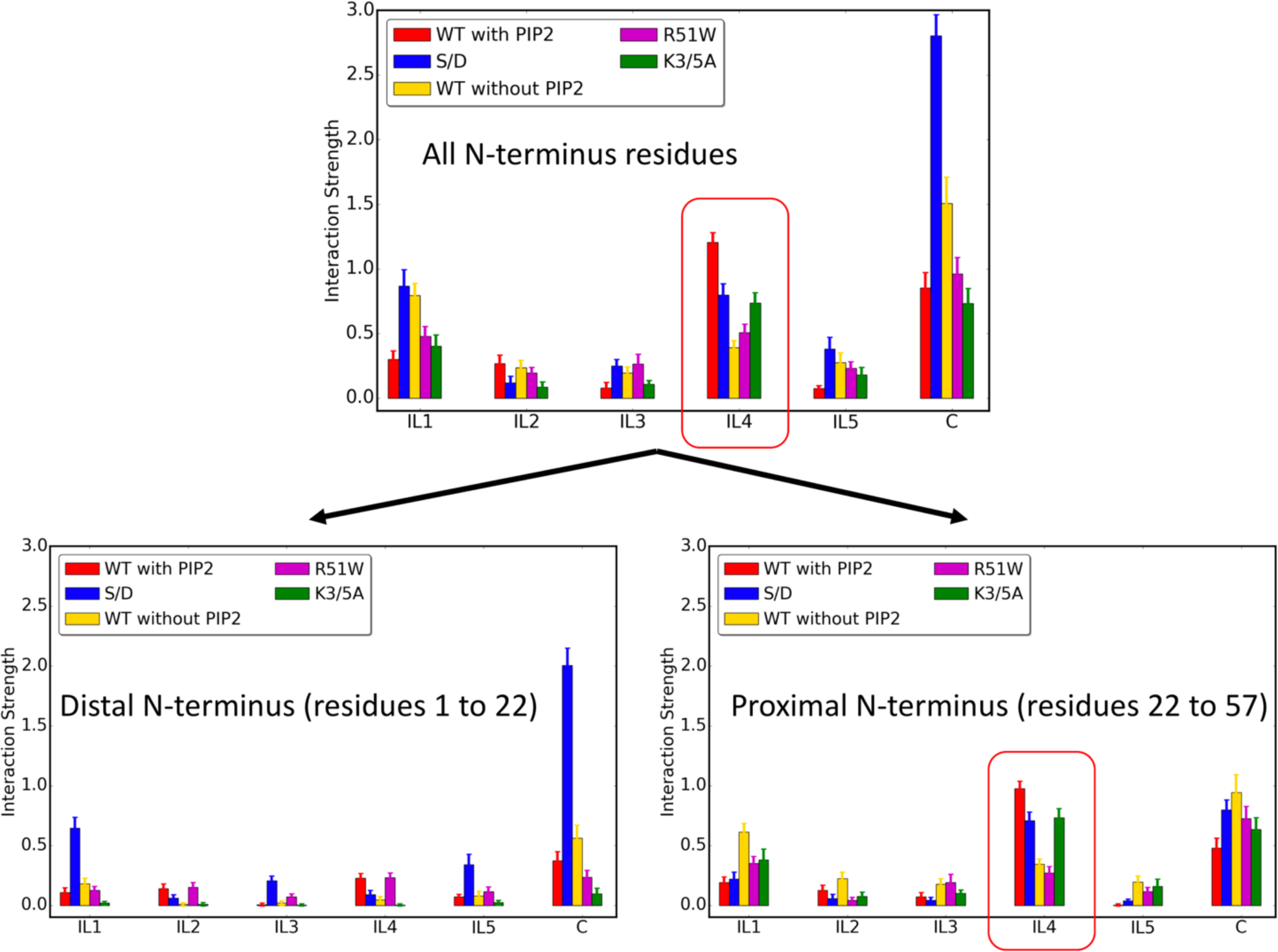
Modes of interaction of hDAT N-terminus with intracellular loop segments. The bars show the average interaction strength calculated from all 50 trajectories of every construct as described in Methods. Error bars show standard deviations (see Methods for full details of calculations and error estimation).

As part of experimental studies of reverse substrate transport (efflux) by DAT (22) and in SERT (34), their N-terminus was truncated, in the case of DAT eliminating the first 22 residues (ΔN22 system). The measurements showed that the truncated transporters maintain direct substrate transport (uptake), but efflux is severely impaired. To compare with and interpret these experimental results, we calculated the modes of interaction of the various constructs for the corresponding components of the N-terminus: the *distal N-terminus* (residues 1 to 22), and the *proximal N-terminus* (residues 23 to 57). Results in Fig. 2 show a distinct difference between the interaction patterns of the two segments. Notably, the largest differences are registered for the interactions of these *distal* and *proximal* segments with IL1, IL4, and the C-terminus. A remarkable similarity is revealed between the <u>pattern</u> of interactions with IL4 calculated for the entire N-terminus, and for just the *proximal* N-terminus (cf. circled regions in Figure 2). This similarity is especially noteworthy because the interaction with IL4 has been singled out to be essential in the early steps of the substrate transport mechanism marked by the release of Na^+^ from the Na2 site (26). Thus, our finding here that the *proximal N-terminus* maintains the essential pattern of interaction with IL4 explains the surprising insensitivity of the inward substrate transport to the deletion of the first 22 residues.

The second largest difference between the interaction patterns of the *proximal* and *distal* N-terminal segments evident in Figure 2 pertains to association with the C-terminus. In particular, the *distal* segment of the phosphomimic S/D construct has a remarkably high interaction quotient with the C-terminus, whereas the weakest association with the C-terminus is observed for the K3/5A construct (in fact, the distal segment of this double mutant K3A/K5A N-terminus is seen to engage in the least amount of interactions with any of the intracellular loops). This diametrically opposed characteristic of the interaction of the two constructs relative to the wild type is remarkable and is fully consistent with the experimentally determined functional properties compared to the wild type hDAT: (i)-only the S/D mutant construct in which the *distal* N-terminus interacts strongly with the C-terminus, has been found thus far to be able to enhance dopamine efflux mediated by DAT in the absence of AMPH (22), (ii)-the K3/5A mutant, which we find to have the weakest interaction between the *distal* segment and the C-terminus, produces a very low amphetamine (AMPH)-induced dopamine efflux (32). The important relation of the *distal* segment with the efflux phenotype is underscored by the deleterious effect of the (ΔN22) truncation on efflux, but not on uptake, as discussed above.

### Multiple paths of inward release of Na^+^ from the Na2 site are regulated by the modes of interaction of the N-terminus

Our detailed study of the release of Na^+^ from the Na2 site (termed Na+/Na2 release) – which is known to initiate solute translocation by the hDAT (28,29) – has identified conformational changes related to function (CCRF), and the underlying allosteric mechanism (30,35,36). These CCRF relate directly to the mode of interaction of the N-terminus with intracellular regions of the transporter in PIP_2_-containing membranes (30). In the earlier studies (26,30) we showed that specific PIP_2_-mediated associations between the N-terminus and various intracellular loop regions of DAT trigger conformational transitions related to the release of Na+/Na2. Here we find, from the new sets of MD simulations of the mutant constructs and conditions we study, that while they differ in their modes of N-terminus interactions (Figure 2), Na+/Na2 release is observed, albeit at different rates, during simulations of the same time length for various (Figures S1 to S4). While the rates of release events observed in the 50 trajectory ensembles for each construct/condition (see Figures S1 to S4) are not rigorously comparable to each other in a statistically meaningful manner, the prediction of inward release of Na+/Na2 in each of these constructs is consistent with experimental results and with our previous finding (30) that the destabilization of the Na+/Na2 is highly correlated with the amount of water penetration to the binding site (Table S4). Figures S5, S6, and S7 show details of the spontaneous release dynamics calculated for the R51W hDAT (Figure S5), the S/D system (Figure S6), and the no-PIP_2_ system (Figure S7). The K3/5A mutant did not exhibit a release event, but the pattern of Na+/Na2 destabilization and intracellular gate opening is similar to that observed for S/D (Figures S3, S4, S8) suggesting that it is on the path to Na+/Na2 release as well.

These results for the large number of different constructs and conditions are remarkably consistent with the experimental evidence showing that the regular transport of the dopamine substrate (uptake) is affected differently by the various mutations/conditions than the reverse transport of this substrate (efflux) induced by amphetamine. Thus, efflux is impaired by most of these mutations/conditions, with the exception of the S/D system which exhibits dopamine efflux even in the absence of AMPH but under elevated intracellular Na^+^ concentrations (25). In particular, experimental evidence points to the importance of PIP_2_-containing membranes for various functional phenotypes of hDAT, including AMPH-induced efflux, but shows that substrate transport is only mildly affected if PIP_2_ content is reduced (32,34,37).

The dependence of functional properties of the eukaryotic transporters on PIP_2_ is not shared by the bacterial transporter homologs, such as the structural prototype LeuT (16,38), which do not require PIP_2_-containing membranes for transport and are also not exhibiting reverse transport. We reasoned that comparing molecular details of functional mechanisms involving the N-terminus in the presence and absence of PIP_2_ would shed new light on the role introduced in evolution by the long N-terminus of the eukaryotic transporters. To discern the source of underlying mechanistic differences that connect PIP_2_ sensitivity to the long N-terminus it became necessary, therefore, to understand (i)-how the initiating step of substrate transport, i.e., the release of Na+/Na2, is achieved in PIP_2_-containing vs PIP_2_-depleted membranes, and (ii)-what the role of the N-terminus interactions is in the CCRF (including Na+/Na2 release process) when PIP_2_ is not present. To this end we used the MSM analysis to obtain a kinetic model for the Na+/Na2 release process in the no-PIP_2_ system, as the comparison of quantitative terms for the wild type protein with/without PIP_2_ allows robust mechanistic inferences as illustrated below.

*Markov State Model analysis of Na+/Na2 release in PIP_2_-depleted membranes:* To enable direct comparison of the results with the MSM analysis of wild type hDAT in PIP_2_-containing membranes (30) we built and analyzed the MSM for the no-PIP_2_ system following the same protocol (see Methods and ref. (30)). Thus, as the same mechanism was followed in the two compared conditions (i.e., with/without PIP_2_), the same set of parameters as before (30) (Table S3) was used to generate the reduced conformational space with the tICA method (see Methods). The tICA energy landscape (Figure 3B) was obtained by projecting all the conformations from all the trajectories onto the first two tICA reaction coordinates. Visualization of conformations belonging to different regions of the tICA energy landscape revealed that this landscape, unlike the one for PIP_2_-containing membrane conditions (shown in Figure 3A), could be divided into only two (rather than three) regions in terms of the location of the Na+/Na2 ion: one in which the Na+/Na2 is still bound in the Na2 site, and the other in which the Na+/Na2 is already released (Figure 3B). Thus, this tICA space (Figure 3B) does not contain a region representing the intermediate state seen in the wild type hDAT system in PIP_2_-enriched membranes when the Na+/Na2 has left the binding site but is not yet released to the intracellular environment because it is interacting with the E428 side chain (Figure 3C). Because this interaction requires the E428 side chain to be free from its partner in the E428–R445 gate (30), the results suggest that a change in N-terminus interactions due to PIP_2_ depletion directly affects this gate. Indeed, the finding summarized in Fig. 2 shows a major reduction in the interactions of the N-terminus with IL4 in the absence of PIP_2_, which is thus seen to result in a more stable R445–E428 gate in the absence of PIP_2_ (Figure 3D, see also Figure S9).

**Figure 3.**
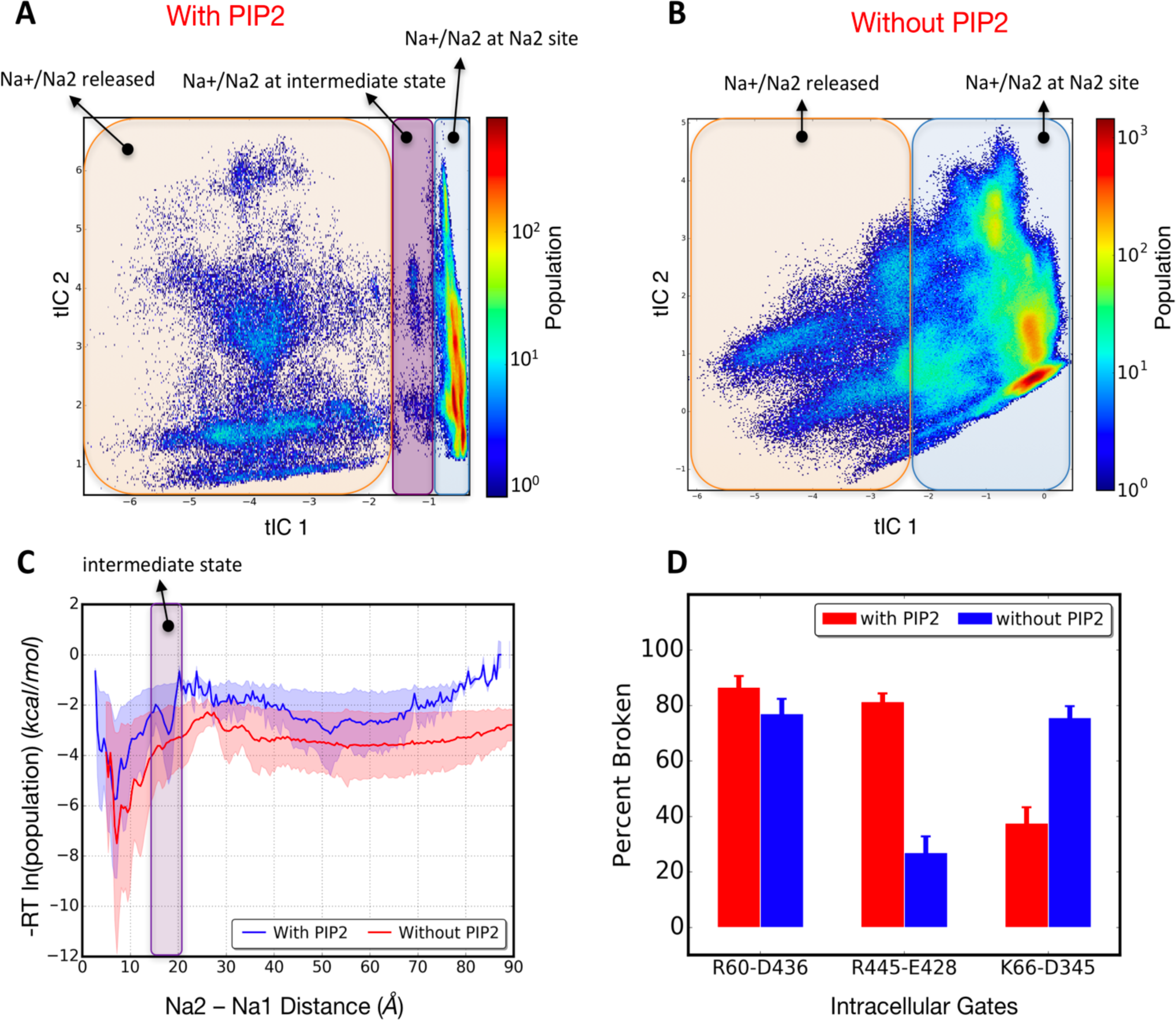
hDAT tICA space in presence and absence of PIP_2_ lipids. (**A** and **B**) Population-weighted tICA landscape for hDAT trajectories in PIP_2_-containing membranes (A) and for the no-PIP_2_ system (B). For each system, all of the conformations in all of the 50 trajectories are projected on the space of the tICA 1^st^ and 2^nd^ eigenvectors. Regions differing with respect to the location of Na+/Na2 are highlighted on the tICA landscape with transparent boxes of different colors. (**C**) All the conformations are projected on the reaction coordinate based on Na+/Na2 distance from the sodium at site Na1, and the free energy (ΔG) is population-based and is calculated as [ΔG=-RT ln(population)]. The intermediate state is highlighted by the magenta box; note the absence of a distinct minimum in the absence of PIP_2_ (red line). Error bars (shown as transparent shades) are calculated using the bootstrap method on 100 blocks of frames with 160 ns time range that are randomly extracted from all 50 trajectories. (**D**) Comparing percent of broken interactions of intracellular gates in the control simulation (PIP_2_-containing membrane) (red bars) and in the no-PIP_2_ system (blue bars) averaged over all 50 trajectories of each construct. Error bars are calculated using the bootstrap method by randomly selecting 50 trajectories (with replacement) and repeating the process for 1000 iterations.

To build the MSM from the 2D tICA landscape shown in Fig. 3B, we followed the same protocol as before (30) to obtain the *implied time scales* plot (see Methods). As shown in Figure S10 (*left panel*), Markovian behavior of the system is observed after a lag time of ~80 ns, therefore the transition probability matrix (TPM) (see Methods) at 80 ns lag time was selected for all subsequent analyses of the no-PIP_2_ system. Mapping all microstates on the tICA landscape and coloring them based on the first MSM eigenvector (i.e., the first TPM eigenvector, shown in Figure S11) reveals that Na+/Na is still bound in microstates with positive sign (red circles in Figure S11), whereas the microstates with negative sign (blue circles in the Figure S11) have released the Na+/Na2 to the intracellular environment. Since the state population flows from the positive to negative states, the first MSM eigenvector is seen to capture the overall release kinetics of the Na+/Na2. The implied time-scale equation (see Methods) shows that this relaxation mode is characterized by time-scales of ~ 1.1 μs, comparable to the previously reported kinetics for hDAT in the PIP_2_-containing membranes (~ 800 ns) (30) (Figure S10).

To compare the mechanisms of sodium release from the Na2 site of the WT hDAT in PIP_2_-containing vs PIP_2_-depleted membranes, we used the same transition path theory (TPT) analysis (see Methods) to obtain the most probable release pathways of Na+/Na2, and quantified the flux associated with each of these on a macrostate–based MSM using 15 macrostates as before (30). Similar to the WT in PIP_2_-containing membranes, several pathways are revealed in the no-PIP_2_ system. Here, the first 10 pathways identified by the TPT analysis contribute ~80% of the total flux between Na+/Na2 bound states and Na+/Na2 released states (highlighted in Figure 4; see Table S5 for quantification of fluxes). Their structural context is shown in Figure 4 and Figure S13.

**Figure 4.**
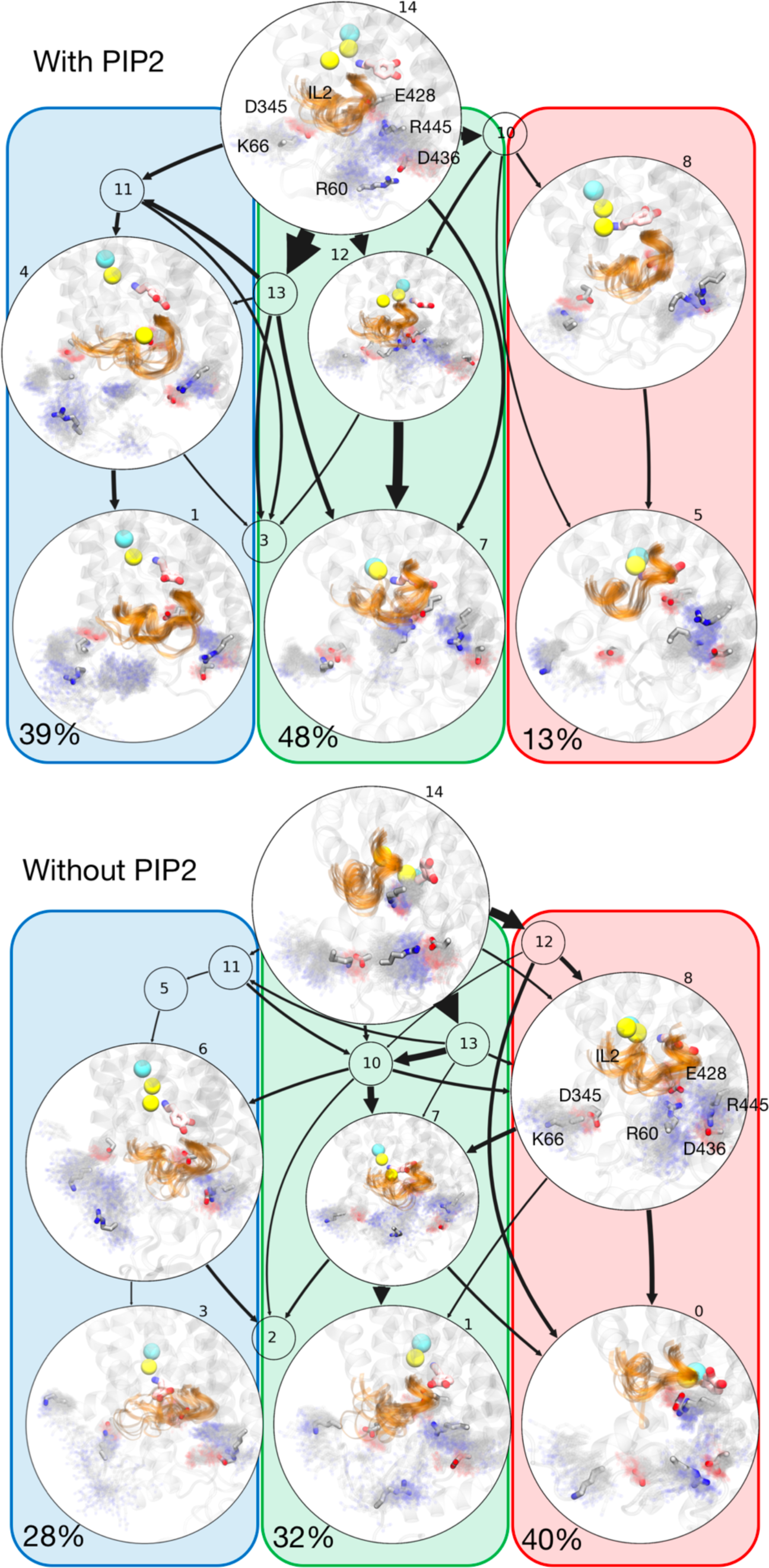
Comparison of Na+/Na2 release pathways calculated in PIP_2_-enriched and PIP_2_-depleted membranes. Results from the transition path theory (TPT) analysis for the release of Na+/Na2 to the intracellular environment by the wild-type hDAT in PIP_2_-containing membranes (top) and in PIP_2_-depleted membranes (bottom) systems. Arrow thickness represents relative flux magnitude for each transition. A representative conformation is shown for selected macrostates, with IL2 highlighted in orange (cartoon representation) on the transparent structure of hDAT. Density representation of the residues forming intracellular gates are shown as fuzzy background gray colors and the most probable locations of these residues within the density representation are highlighted in licorice rendering and labeled for reference. Dopamine is shown in pink licorice and sodium and chloride ions in yellow and cyan spheres, respectively. Red, green, and blue boxes highlight the three major release pathways of Na+/Na2; numbers indicate the fraction of the total flux carried by each major pathway.

The identification of the major Na+/Na2 release pathway in the no-PIP_2_ system reveals a clear difference from the release in the PIP_2_-enriched system (cf. panels labeled “with PIP_2_” and “without PIP_2_” in Figure 4) that echoes the difference in the modes of interaction described by the results in Fig. 2. In the no-PIP_2_ system, the first major pathway for the release contributes ~40% of the total flux and is formed when both the R60–D436 and the E428–R445 gates remain closed (a consequence of the much lower interaction with IL4 as seen in Fig. 2), but the K66–D345 is open (highlighted in the transparent red box in Figure 4). This is in sharp contrast to the result in the presence of PIP_2_ where this pathway is a minor contributor to the total flux, at only ~13%.

The second major pathway in the no-PIP_2_ system is enabled when both R60–D436 and E428–R445 gates are broken, but R60 is now able to form interactions with E428 (Figure 4, green box). This pathway contributes ~32% to the total flux. Notably, this was the major Na+/Na2 release pathway in the presence of PIP_2_, contributing about 48% percent of the total flux. But because in the absence of PIP_2_ R60 (or R445) is interacting with E428, the intermediate state cannot form.

The third major pathway in the no-PIP_2_ system is formed when the R60–D436 gate is broken and the N-terminus moves closer to IL2. The intracellular gate interaction E428–R445 is still maintained (Figure 4, blue box). In the presence of PIP_2_, this pathway contributes about ~39% to the total flux, but in the no-PIP_2_ system the contribution is reduced to 28%. Importantly, a major difference is that in the presence of PIP_2_, R60 is seen to be engaged in PIP_2_-mediated interactions with several positively charged residues from IL2 (K257, K260, K264), whereas in the no-PIP_2_ system the association between the N-terminus and the IL2 region still permits interactions between R60 and D68 and/or D385 (see macrostate 3 in Fig. 4 and Fig. S12).

Overall, the TPT analysis shows that the preference ranking of the various release pathways, which we quantified by calculating fluxes, is strongly affected by the PIP_2_ lipids in a manner consistent with the modes of interaction with the N-terminus. The difference in Na+/Na2 mechanism between the two conditions (with/without PIP_2_) is underscored by the finding that the increased dynamics of the N-terminus (Figure S12) in the absence of PIP_2_ results in a destabilization of the K66–D345 gate, which enhances the Na+/Na2 release flux through a pathway that was only marginally active in the presence of PIP_2_ (pathway highlighted in red box). Taken together, these results show how the eukaryotic transporters can adapt to different membrane composition conditions by utilizing different N-terminus interaction patterns so that release of the Na^+^ from the Na2 site is maintained.

## Discussion

The extensive investigations of the molecular mechanisms underlying the vital role of human DAT in signal transduction have profited much from the availability of structurally simpler prototypes of the NSS, such as the bacterial analog LeuT (a leucine and alanine transporter), for which the crystallographic data provided the first structural basis for detailed molecular studies (see (16,38)). When the striking fold-similarity of LeuT to the eukaryotic and human neurotransmitter transporters such as DAT was established (39,40), it enabled the major progress in understanding the functional mechanism documented in a very large number of publications (for reviews see (41)). It became clear, however, that various physiologically important mechanisms that eukaryotic transporters such as DAT and SERT have acquired through evolution involve allosteric coupling to their environment that differentiate them from the structurally simpler bacterial analogs. In a large number of studies (20,22,24-26,30,32,42-51) to which we have contributed findings from both computation and experiments, these new functions of the eukaryotic transporters – such as regulation by lipids, by phosphorylation, and the observed efflux phenotype – were shown to be mechanistically modulated by defined structural elements, especially the relatively long N- and C-terminal segments that are found exclusively in this class, and not in bacterial counterparts (40). These intracellularly located N- and C-terminal segments are the most divergent structural units within the NSS family of proteins, as they range in size from just a few (~10) amino acids (in bacterial members, such as LeuT) to segments containing > 60 residues (N-terminus of eukaryotic NSS such as DAT, SERT, see (40,52)).

To understand how the new functions of the eukaryotic transporters relate to the presence of the N-terminus and its dynamic properties, we quantified the interaction of the hDAT N-terminus with the intracellular regions of the hDAT transporter. On this basis we were able to relate different modes of interaction to the functional mechanisms of DAT through the relation between the experimentally measured effects of mutations/conditions, and the changes in modes of interaction. The release of the Na^+^ ion from the Na2 site, which is known to be a key first step in the substrate translocation cycle (28,29), served as an established functional readout for the initiation of the transport cycle. We showed how the effects of modes of interaction of the N-terminus with various intracellular regions of the transporter relate to various modes of intracellular gate opening and paths of water penetration. This led to a mechanistic interpretation of the experimentally measured modifications of the functional properties observed for mutant constructs in the presence and absence of PIP_2_. Together, our results reveal how changes in the modes of interaction stemming from N-terminus mutations and PIP_2_ depletion, are directly associated with stabilization/destabilization of intracellular gates, and their effect on the penetration of water into the binding site (Figure 5A) that is required for uptake and efflux in hDAT.

**Figure 5.**
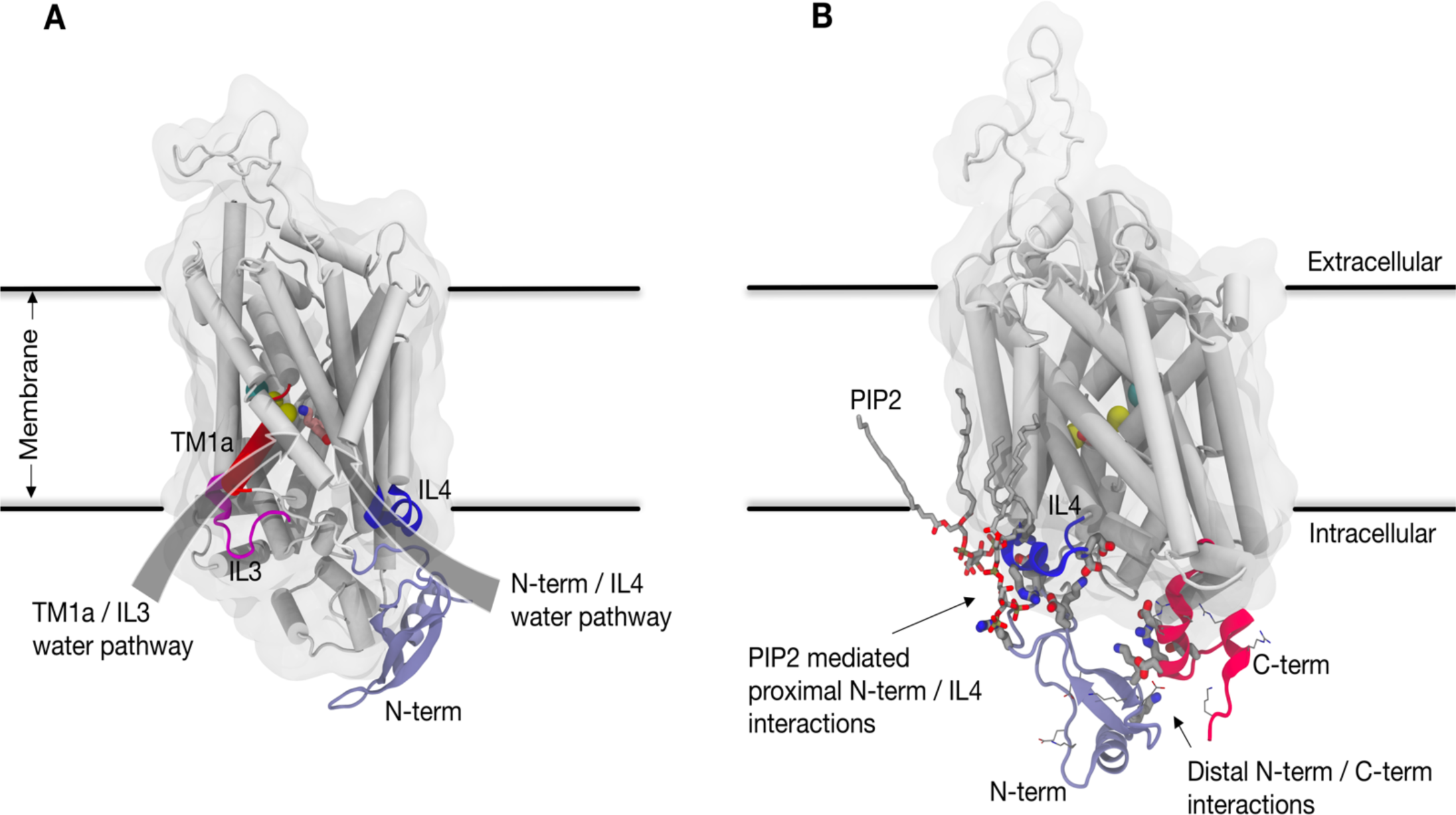
Illustration of effects of different modes of N-terminus interactions on functional phenotypes. (**A**) Representation of two water penetration pathways from the intracellular side of the hDAT. The N-terminus/IL4 water pathway is the main water penetration path in the wildtype in the presence of PIP_2_ lipids. In the absence of PIP_2_, the N-terminus/IL4 water (blue) penetration pathway is impaired, and hydration of the binding site is achieved primarily through another water pathway formed between TM1a and IL3 (red). (**B**) Representation of conditions linked to the efflux function of hDAT. The interaction of the proximal N-terminus with IL4 (blue) is mediated by PIP_2_ lipids, and the distal part of the N-terminus engages with the C-terminus (red). All charged residues are shown for both N-terminus and C-terminus with thin licorice; charged residues engaged in interactions between N-terminus and either IL4 or the C-terminus, are highlighted with thick licorice. Dopamine is shown in the S1 binding site (in pink), sodium ions are shown as yellow spheres, and the chloride ion is in cyan.

By specifying the modes of interactions of the N-terminus that regulate the opening probabilities of water pathways separately for the *distal* and the *proximal* segments of the N-terminus, our findings provide a clear mechanistic explanation for a persistent conundrum in the literature regarding the effects of (i)-truncating the first 22 residues of the N-terminus, and (ii)-PIP_2_ depletion, and in particular their different effects on the uptake and efflux aspects of eukaryotic transporter function. That the truncation has little effect on uptake is explained by our finding (summarized in Figure 2) that the pattern of interactions of the N-terminus with IL4 of DAT is mimicked by just the *proximal* segment, without special involvement of the *distal* segment. However, this interaction that has been shown to disrupt the E428–R445 gate and thus serve as one of the major mechanistic triggers for Na+/Na2 release, is PIP_2_-mediated (26), and is reduced under no-PIP_2_ conditions (Figure 2). This might have suggested a reduced release of Na+/Na2 that would impair the substrate uptake function under these conditions. Yet this is not what is observed experimentally, as the uptake function of hDAT is not impaired by PIP_2_ depletion from the membrane. The mechanistic explanation emerges from our analysis of (a)-the simulations presented here that show how modes of interaction of the N-terminus change under the different conditions (and for the various mutant constructs; Figure 2), and (b)-the MSM from the simulation trajectories of the no-PIP_2_ condition showing how Na+/Na2 release is maintained, but with different probabilities of the release pathways that involve alternative water penetration channels (e.g., the TM1a/IL3 channel (Figure 5A) that is formed when K66–D345 interaction breaks).

The effect of N-terminus truncation on efflux is even more interesting in the context of the functional difference between the bacterial and eukaryotic transporters because elimination of the *distal* segment in the neurotransmitter SLC6 transporters impairs a function not shared with the bacterial homologs. The relation of the *distal* segment to efflux makes it tempting to speculate that some combination of the interaction modes of the *proximal* N-terminus (Figure 5B) is required for efflux. Indeed, the results summarized in Figure 2 show (i)-a reduction of *proximal* N-terminus/IL4 interactions in constructs with low efflux activity (R51W and K3/5A) and in PIP_2_-depleted membranes, and (ii)-a reduction in *distal* N-terminus/C-terminus interactions in the efflux-deficient K3/5A mutant, and an increase of these interactions in the efflux-promoting S/D construct. An involvement of the *distal* N-terminus/C-terminus interactions in facilitating efflux is also consistent with the observation that palmitoylation of the C-terminus reduces efflux, presumably by limiting interaction with the C-terminus. This limitation would reduce N-terminus phosphorylation (53,54) by the Ca2+/calmodulin-dependent protein kinase II (CaMKII) that is proposed to attach to the distal C-terminus of hDAT in order to phosphorylate the serine residues in the distal N-terminus for AMPH-induced efflux (25).

The mechanistic explanations for the experimental findings about the modulation of function by the N-terminus of DAT, SERT, and other eukaryotic transporters, connect the elongation of N- and C-termini in the evolution from bacterial homologs, with the appearance of new functional properties (e.g., efflux). These functional phenotypes, enabled by specific roles of the *proximal* and *distal* segments, are not shared by the bacterial homologs which lack the long N-terminus but share the overall molecular architecture (termed the “LeuT-fold”; (38)). The specific (different) involvement of the *proximal* and *distal* segments – such as the role of the *proximal* segment in sustaining transport in PIP_2_-depleted membranes, and of the *distal* segment in modulating efflux – may represent an evolutionary adaptation required for the function of eukaryotic transporters expressed in various cell types of the same organism that differ in the lipid composition and protein complement of the membrane environment.

## Methods

### System preparation

The molecular model of full-length wild type hDAT used in this study is the same as described and investigated earlier (30). The R51W and K3A+K5A constructs were prepared by introducing the mutations in the wild type hDAT model using VMD mutator plugin (55). To build the S/D mutant (simultaneous mutations of S2, S4, S7, S12, and S13 residues to Asp), we combined, using Modeller version 9v1 (56), the 57-620 residue stretch from the wild type hDAT structure with the structural model of the 1-57 S/D segment elaborated and described previously (20).

The full length models of the hDAT constructs R51W, K3A+K5A (heretofore referred to as “K3/5A”), and S/D were inserted into the same pre-equilibrated compositionally asymmetric bilayer membrane used for MD simulations of the wild type hDAT (30). This lipid bilayer was designed to mimic a neuronal cell plasma membrane and contains 5% PI(4,5)P_2_ lipid on the intracellular leaflet of the bilayer (see Table S1 for membrane lipid composition). For the simulations of the wild type hDAT in PI(4,5)P_2_-depleted membrane environment, as done previously (26), all the PI(4,5)P_2_ lipids in the bilayer were changed to POPE lipids, the major component of the intracellular leaflet of our model bilayer. All the hDAT-membrane systems were solvated in a 150 mM K+Cl^-^ TIP3P water solution with ions added for neutrality, resulting in a final atom count of ~150,000.

### Molecular dynamics simulations

All-atom molecular dynamics (MD) simulations were carried out using the same scheme as described earlier for the wild type hDAT in PI(4,5)P_2_-enriched membranes (30). Briefly, using NAMD software version 2.10 (57), the systems first were equilibrated following the same multi-step equilibration protocol used previously (30) during which the backbone of the protein was first fixed, then harmonically restrained, and finally released. After this equilibration phase, the velocities of all the atoms in the system were reset (at T=310K using random number seed) and 50 independent ~1μs long unbiased MD simulations were carried out using the latest version of the ACEMD software (58) resulting in a cumulative MD simulation time of ~50 μs per system. These production simulations were performed under NVT ensemble and with all the default run parameters validated by the ACEMD developers (https://www.acellera.com/) and in a large number of published applications (e.g., see https://www.acellera.com/products/molecular-dynamics-software-GPU-acemd/use-cases/). The run parameters (4fs time step with hydrogen mass repartitioning; PME for electrostatics; switched Lennard-Jones interactions with a cut-off of 9Ǻ, and switching distance set to 7.5Ǻ) have been shown to reliably reproduce known values for free energy of protein folding and a variety of properties of lipid membranes (59,60). In addition, ensemble MD simulations with ACEMD have been generally used to generate large data sets of trajectories for quantitative analysis of kinetics of ligand-induced conformational transitions in G protein-coupled receptors (GPCRs) (61), of protein-protein association/dissociation processes (62), of phospholipid scrambling process mediated by the GPCR opsin (63) as well as for identifying pathways for spontaneous cholesterol movement in adenosine A2A GPCR (64).

### Calculation of the interaction strengths

To obtain a measure of interaction between N-terminus and other intracellular regions of hDAT we counted the number of interactions between charged residues from the N-terminus and the intracellular loop regions. The number of interactions was quantified for a cutoff distance of 7 Å between interacting residue pairs from head group atoms of the N-terminus and of intracellular domain residues (using N_ξ_ for Lys, C_ξ_ for Arg, C_δ_ for Glu, and C_γ_ for Asp). The residues included in the calculations are listed in Table S2. All 50 trajectories for each construct were used for these calculations, with the first 500 ns of each trajectory considered an equilibration phase so that only the 500-940 ns time interval from each trajectory was used to ensure that the total number of frames used for the analysis is the same for each construct. Total number of interactions is then divided by total number of trajectories (i.e. 50) to obtain the average “interaction strength” represented in Figure 2. Error bars are calculated by taking the standard deviation of “interaction strength” in all 50 trajectories.

### Markov State based quantitative kinetic model construction

We used the Markov State Model (MSM) approach to analyze the trajectories in the absence of PIP_2_ lipids and build quantitative kinetic models of sodium release from the Na2 site for comparison with the kinetics and pathways calculated in the presence of PIP_2_ in Ref. (30). Therefore, we have followed here the same protocol as described in detail in Ref. (30). Such quantitative kinetic models provided by Markov State Models (MSMs) (65,66),(67) have been widely applied in protein folding studies (68,69) and MSM based kinetic models predictions have been validated experimentally (70,71). We and others have used MSMs combined with reactive flux analysis, such as transition path theory (TPT) analysis to obtain key mechanistic insights into functions of membrane proteins (30,72,73). The three main components for achieving quantitative MSM-based kinetic models are briefly reviewed below.

*1. <u>Dimensionality reduction using tICA</u>*. Reducing the dimensionality of a system as large and complex as the membrane-immersed hDAT is necessary in order to construct an interpretable kinetic model. A natural choice of suitable reaction coordinates is for those that can project the conformational space of the system along its *slowest* reaction coordinate, as this reaction coordinate will capture most of the conformational heterogeneity during the time-course of the simulation (74). The *time-structure based Independent Component Analysis* (tICA) method was developed recently for this purpose of capturing the slowest reaction coordinate of a system (75-77). Briefly, the tICA method involves a transformation that utilizes two matrices constructed from the trajectory data: the covariance matrix **C**, and a time-lagged covariance matrix ***C_TL_***. The slowest reaction coordinates of a system are then defined by eigenvectors of the generalized eigenvalue problem: ***C_TL_*V** = **CVΛ**, where **Λ** and **V** are the eigenvalue and eigenvector matrices, respectively. The eigenvectors corresponding to the largest eigenvalues identify the slowest reaction coordinates. Here we used a lag time of 16 ns to construct the time-lagged covariance matrix ***CTL***, and the tICA parameters were chosen as before for the hDAT molecular system (30), to measure a) the dynamics of the Na^+^ ion from the Na2 site, termed Na+/Na2, and b) the dynamics of intracellular gates formed between residues R60, D436, R445, E428 (Table S3).

*2. <u>Markov Model construction.</u>* The conformational transitions of biomolecular systems where the time between transitions is long enough can be modeled as Markov chains (65) in which transitions among states depend only on the current state of the system (i.e., Markovian behavior). Such Markov models provide powerful tools for outcome prediction by enabling the extraction of long time-scale information from multiples of short time-scale events.

Two components needed for the construction of such a Markov model are: an ensemble of *microstates* of the system, and of the transitions among these microstates (78). Microstates are defined by clustering the conformational space of the system into several basins using automated clustering algorithms like *K*-means or *K*-centers, and is most practical if performed in a dimensionality-reduced space such as the one obtained from the tICA transformation. The transitions among the microstates are calculated for a particular time interval between each of the transitions (so called lag-time) and stored in the *transition count matrix*. By row-normalizing the transition count matrix one obtains the *transition probability matrix* (TPM). To validate Markovian behavior, the TPMs are constructed for multiple lag times and the relaxation timescales of the system are extracted by using the relation:

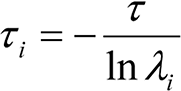

where *t* is the lag-time used for building the TPM, λ_i_ is the *i*^th^ eigenvalue of the transition probability matrix, and the resulting *τ_i_* is the so-called *implied timescale* corresponding to the *i*^th^ relaxation mode of the system. The system is considered to be Markovian if the behavior is such that *τ_i_* is independent of *τ*′; the corresponding TPM is a *Markovian* TPM that contains all the information about the thermodynamics and kinetics of the system. Thermodynamic information is stored in the first eigenvector (which has an eigenvalue of 1). Information about kinetics is stored in the subsequent (second, third, etc.) eigenvectors, with the second eigenvector (corresponding to the second largest eigenvalue) capturing the slowest dynamics of the system, the third eigenvector capturing the second slowest dynamics, and so on.

Following the same protocols as described in detail in Ref. (30) for the construction of the Markov Models we discretized the reduced conformational space generated by the first two tICA reactions coordinates into 100 microstates (Figure S11) using the *K-means* clustering algorithm implemented in the MSMBuilder3 software (79). Transition probability matrices (TPMs) were constructed at several different lag times to obtain the implied time-scale plots shown in Figures S10, so that the one in which Markovian behavior is exhibited can be identified and chosen for further analysis.

*3. <u>Transition path theory analysis</u>*. In addition to the thermodynamics and kinetics information it contains, the Markovian TPM also contains mechanistic information for a dynamic system. An established means of revealing such mechanistic information inherent in the TPM is the Transition Path Theory (TPT) analysis that identifies the most probable flux pathways of the system (80). TPT provides such pathways by constructing a *flux matrix* from the Markovian TPM. This matrix conversion has been documented in detail (80,81) and its implementation is discussed in our previous publication (30). Although directly applicable to MSM in the microstate space (on the order of hundreds to thousands of states), TPT analysis is usually done on a *macrostate* MSM (on the order of tens of states) for a better visualization of flux pathways. Here we transitioned from the microstate–MSM to macrostate–MSM by using the Robust Perron Cluster Analysis (PCCA^+^) algorithm (82) that lumps microstates into macrostates using the sign structure of the MSM eigenvectors (this assumes that microstates with the same signs, positive or negative, in the MSM eigenvectors, will have similar kinetics (83)). Using the PCCA^+^ algorithm we obtained 15 macrostates and by applying the TPT analysis to these macrostates we obtained the most probable flux pathways for the system.

## List of abbreviations

SLC6: solute carrier 6
NSS: neurotransmitter:sodium symporter
DAT: dopamine transporter
hDAT: human dopamine transporter
SERT: serotonin transporter
TM: transmembrane
MSM: Markov State Model
tICA: time-structure based independent component analysis
TPM: transition probability matrix
TPT: transition path theory

## Declarations

**Competing interests:** The authors declare that they have no competing interests.

**Funding**: The work was supported by NIH Grants P01 DA012408, R01 DA041510 and in part U54 GM087519.

**Authors’ contributions:** All authors contributed to conceiving the project and experimental design, A.M.R. did the analysis and carried out the MSM construction, G.K. carried out the MD simulations, all authors participated in the interpretation of results and in writing the manuscript.

## Acknowledgments

The following computational resources are gratefully acknowledged: resources of the Oak Ridge Leadership Computing Facility (ALCC allocation BIP109) at the Oak Ridge National Laboratory, which is supported by the Office of Science of the U.S. Department of Energy under Contract No. DE-AC05-00OR22725; an allocation at the National Energy Research Scientific Computing Center (NERSC, repository m1710) supported by the Office of Science of the U.S. Department of Energy under Contract No. DE-AC02-05CH11231; and the computational resources of the David A. Cofrin Center for Biomedical Information in the HRH Prince Alwaleed Bin Talal Bin Abdulaziz Alsaud Institute for Computational Biomedicine at Weill Cornell Medical College.

